# Benchmarking artificial intelligence methods for end-to-end computational pathology

**DOI:** 10.1101/2021.08.09.455633

**Authors:** Narmin Ghaffari Laleh, Hannah Sophie Muti, Chiara Maria Lavinia Loeffler, Amelie Echle, Oliver Lester Saldanha, Faisal Mahmood, Ming Y. Lu, Christian Trautwein, Rupert Langer, Bastian Dislich, Roman D. Buelow, Heike Irmgard Grabsch, Hermann Brenner, Jenny Chang-Claude, Elizabeth Alwers, Titus J. Brinker, Firas Khader, Daniel Truhn, Nadine T. Gaisa, Peter Boor, Michael Hoffmeister, Volkmar Schulz, Jakob Nikolas Kather

## Abstract

Artificial intelligence (AI) can extract subtle visual information from digitized histopathology slides and yield scientific insight on genotype-phenotype interactions as well as clinically actionable recommendations. Classical weakly supervised pipelines use an end-to-end approach with residual neural networks (ResNets), modern convolutional neural networks such as EfficientNet, or non-convolutional architectures such as vision transformers (ViT). In addition, multiple-instance learning (MIL) and clustering-constrained attention MIL (CLAM) are being used for pathology image analysis. However, it is unclear how these different approaches perform relative to each other. Here, we implement and systematically compare all five methods in six clinically relevant end-to-end prediction tasks using data from N=4848 patients with rigorous external validation. We show that histological tumor subtyping of renal cell carcinoma is an easy task which approaches successfully solved with an area under the receiver operating curve (AUROC) of above 0.9 without any significant differences between approaches. In contrast, we report significant performance differences for mutation prediction in colorectal, gastric and bladder cancer. Weakly supervised ResNet-and ViT-based workflows significantly outperformed other methods, in particular MIL and CLAM for mutation prediction. As a reason for this higher performance we identify the ability of ResNet and ViT to assign high prediction scores to highly informative image regions with plausible histopathological image features. We make all source codes publicly available at https://github.com/KatherLab/HIA, allowing easy application of all methods on any end-to-end problem in computational pathology.

## Introduction

Artificial intelligence (AI) methods are widely used for end-to-end analysis of histopathological whole slide images (WSI) in the research platform. Classical applications of such end-to-end workflows are tumor detection [1], subtyping [2,3] and grading [4] which can recapitulate, automate and improve pathologists’ assessment of WSI. However, AI has also been used to perform image analysis tasks which exceed human capabilities, including prediction of molecular alterations [5], prognostication [6] and prediction of treatment response [7] directly from routine WSI. Collectively, these broad applications of AI in WSI image analysis are termed “computational pathology” and widespread clinical adoption is ultimately expected once routine diagnostic workflows are fully digitalized. [8] As glass slides stained with hematoxylin and eosin (H&E) are ubiquitously available for almost every cancer patient, uptake of AI methods in clinical routine is expected to integrate in existing diagnostic pathways, improve outcomes and provide cost savings. [9]

However, a major limitation for the development, validation and commercialization of computational pathology methods is the lack of systematic comparison (i.e. benchmarking) of different technologies. While the earliest studies in 2018 employed a weakly-supervised approach based on a convolutional neural network (CNN) and spatial averaging [10], recent studies have proposed conceptually new technologies, including attention-based methods [11] and multiple-instance learning [1,2,12]. In addition, computational pathology is an applied field which follows trends in basic computer vision research. Thus, it can be anticipated that classical CNN architectures such as ResNets (Residual Neural network) will be ultimately replaced by more powerful and efficient CNNs such as EfficientNet [13] or non-convolutional AI approaches such as Vision Transformers (ViT) [14]. However, for academic and commercial actors in the field of computational pathology, choosing the best method for an end-to-end problem is currently not possible on a conceptual and practical level. On a conceptual level, there is currently no systematic evidence on which methods yield the best performance for clinically relevant problems, preventing researchers, pathologists and companies from making optimal design choices for a computational pathology application. On a practical level, there is currently no implementation of the whole spectrum of AI methods for computational pathology.

In the present study, we systematically collected WSI datasets for six clinically common end-to-end prediction tasks with diagnostic or therapeutic relevance. In renal cell carcinoma, we investigated the classification of morphological subtypes, which is a widely studied problem [2]. In colorectal cancer, we investigated AI-based prediction of the immunotherapy biomarker microsatellite instability (MSI) [15] and mutations in the BRAF gene, which is a directly targetable genetic alteration [16]. In gastric cancer, we investigated prediction of established or potential biomarkers for immunotherapy MSI and Epstein-Barr virus (EBV) positivity. Finally, in bladder cancer, we investigated prediction of FGFR3 mutational status, which is a clinically approved therapeutic target. [17] For each of these tasks, we presented datasets from two different institutions, allowing us to benchmark all different AI approaches with external validation (**Figure 1 A-E**).

**Figure 1.**
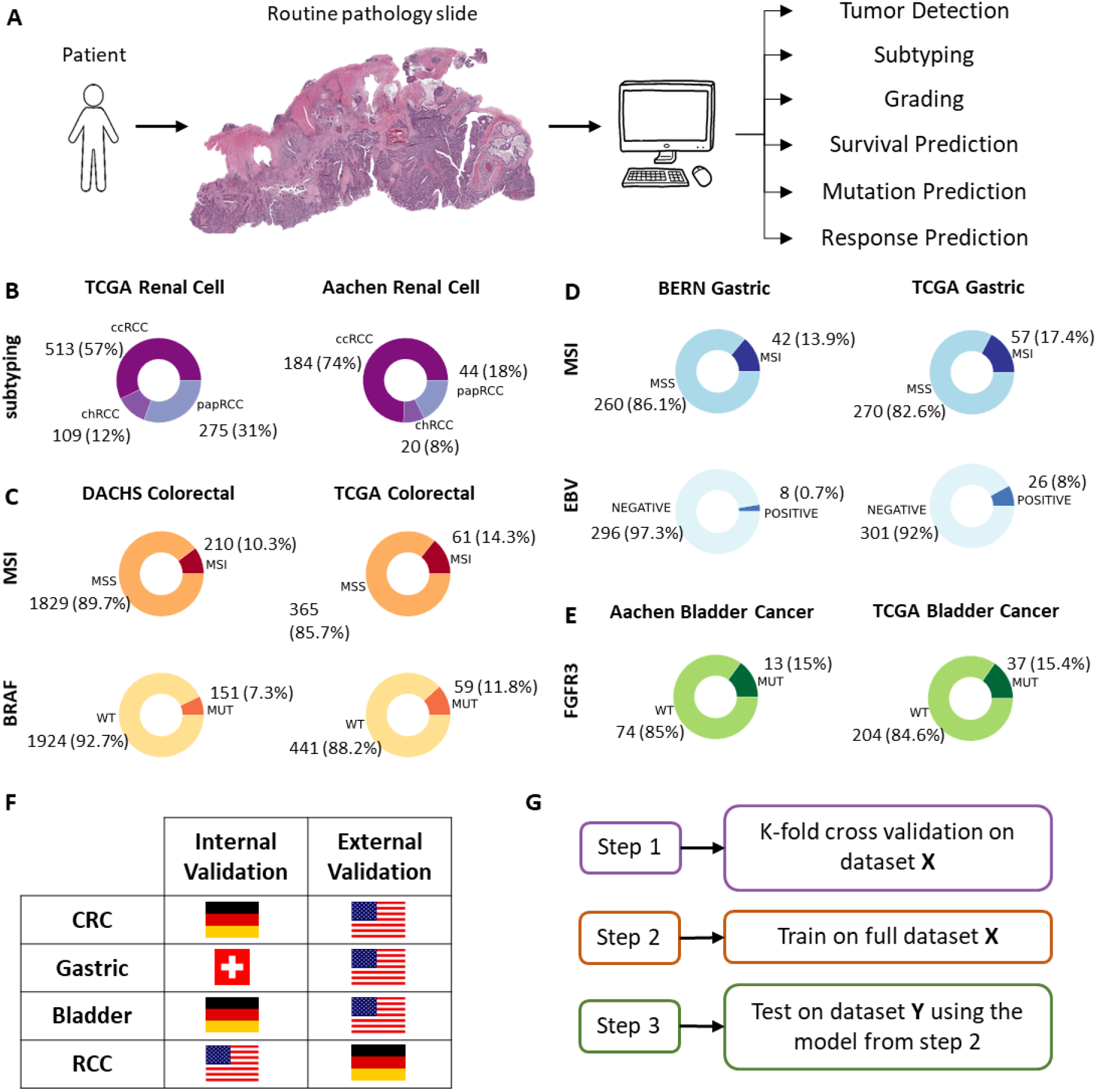
Outline of this study. A) End-to-end artificial intelligence (AI) methods in computational pathology are used to predict a range of features. B) Patient cohorts for renal cell carcinoma, C) for colorectal cancer, D) for gastric cancer and E) for bladder cancer. F) Country of origin of all cohorts. G) Experimental design in this study.

## Results

### All methods achieve high performance for subtyping of renal cell carcinoma

Morphological subtyping of renal cell carcinoma (RCC) into clear cell, chromophobe and papillary subtypes is a widely-studied and clinically relevant problem. Using the “The Cancer Genome Atlas” (TCGA) cohort (TCGA-RCC, N=897 patients, **Suppl. Table 1**), we benchmarked classification performance of end-to-end prediction workflows based on ResNet, EfficientNet and ViT as well as classical MIL and clustering-constrained attention MIL (CLAM, **Figure 2**). We found that in a stratified three-fold cross validation, all methods achieved a high classification performance with macro-averaged area under the receiver operating curve (AUROC) values above 0.90 (**Table 1, Figure 3 A-C**). ViT achieved the highest absolute performance with AUROCs of 0.984 (with 90% confidence interval of 0.977 -0.991), 0.993 (0.988 -0.997) and 0.988 (0.984 -0.993) for detection of all three classes. The classical ResNet-based approach yielded AUROCs of 0.978 (0.970 -0.985), 0.986 (0.980 -0.991) and 0.984 (0.976 -0.992), demonstrating the efficiency of simple classical methods. While MIL-based methods yielded a high absolute performance, this was consistently lowest in all target classes, with classical MIL achieving AUROCs of 0.961 (0.947 -0.973), 0.961 (0.947 -0.972) and 0.957 (0.932 -0.977). Next, we trained classifiers on all TCGA cases and validated them on our in-house dataset (N=248 patients). As expected, performance values slightly decreased, but ViT remained the highest-scoring approach with AUROCs of 0.973 (0.958 -0.985), 0.971 (0.929 -0.998) and 0.97 (0.952 -0.984) for all classes (**Table 2**) and high areas under the precision recall curve (AUPRCs, **Suppl. Table 2**). However, the performance differences between all methods compared to Resnet (**Suppl. Table 3**), EfficientNet (**Suppl. Table 4**), ViT (**Suppl. Table 5**), MIL (**Suppl. Table 6**) and CLAM (**Suppl. Table 7**) did not reach statistical significance in the external validation experiments. We conclude that AI-based RCC subtyping is achievable with almost perfect accuracy compared to the ground truth by any of the tested computational pathology methods.

**Table 1:**
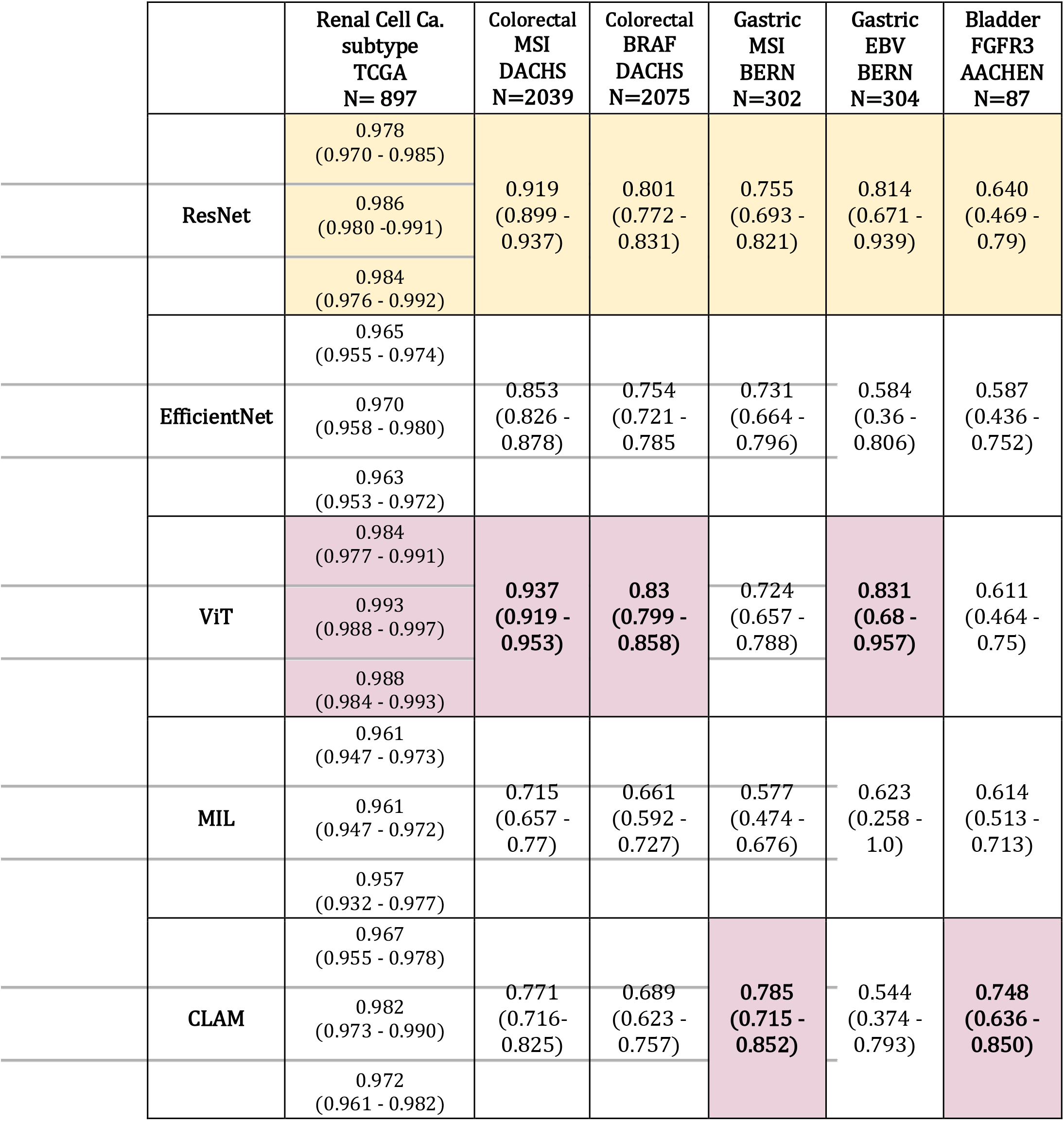
Performance statistics for within-cohort experiments. Performance was assessed by stratified three-fold patient-level cross-validation. Performance is reported as patient-level area under the receiver operating curve (AUROC) with a 90% confidence interval obtained by 1000x bootstrapping. Pink = best, yellow = second-best. For RCC subtyping, AUROCs from top to bottom refer to clear cell, chromophobe and papillary RCC.

|  | Renal Cell Ca.<br>subtype<br>TCGA<br>N= 897 | Colorectal<br>MSI<br>DACHS<br>N=2039 | Colorectal<br>BRAF<br>DACHS<br>N=2075 | Gastric<br>MSI<br>BERN<br>N=302 | Gastric<br>EBV<br>BERN<br>N=304 | Bladder<br>FGFR3<br>AACHEN<br>N=87 |
| --- | --- | --- | --- | --- | --- | --- |
|  | 0.978<br>(0.970 - 0.985) |  |  |  |  |  |
| **ResNet** | 0.986<br>(0.980 - 0.991) | 0.919<br>(0.899 - 0.937) | 0.801<br>(0.772 - 0.831) | 0.755<br>(0.693 - 0.821) | 0.814<br>(0.671 - 0.939) | 0.640<br>(0.469 - 0.79) |
|  | 0.984<br>(0.976 - 0.992) |  |  |  |  |  |
|  | 0.965<br>(0.955 - 0.974) |  |  |  |  |  |
| **EfficientNet** | 0.970<br>(0.958 - 0.980) | 0.853<br>(0.826 - 0.878) | 0.754<br>(0.721 - 0.785) | 0.731<br>(0.664 - 0.796) | 0.584<br>(0.36 - 0.806) | 0.587<br>(0.436 - 0.752) |
|  | 0.963<br>(0.953 - 0.972) |  |  |  |  |  |
|  | 0.984<br>(0.977 - 0.991) |  |  |  |  |  |
| **ViT** | 0.993<br>(0.988 - 0.997) | 0.937<br>(0.919 - 0.953) | 0.83<br>(0.799 - 0.858) | 0.724<br>(0.657 - 0.788) | 0.831<br>(0.68 - 0.957) | 0.611<br>(0.464 - 0.75) |
|  | 0.988<br>(0.984 - 0.993) |  |  |  |  |  |
|  | 0.961<br>(0.947 - 0.973) |  |  |  |  |  |
| **MIL** | 0.961<br>(0.947 - 0.972) | 0.715<br>(0.657 - 0.77) | 0.661<br>(0.592 - 0.727) | 0.577<br>(0.474 - 0.676) | 0.623<br>(0.258 - 1.0) | 0.614<br>(0.513 - 0.713) |
|  | 0.957<br>(0.932 - 0.977) |  |  |  |  |  |
|  | 0.967<br>(0.955 - 0.978) |  |  |  |  |  |
| **CLAM** | 0.982<br>(0.973 - 0.990) | 0.771<br>(0.716 - 0.825) | 0.689<br>(0.623 - 0.757) | 0.785<br>(0.715 - 0.852) | 0.544<br>(0.374 - 0.793) | 0.748<br>(0.636 - 0.850) |
|  | 0.972<br>(0.961 - 0.982) |  |  |  |  |  |

**Table 2:** Performance statistics for external validation experiments. Performance is reported as patient-level area under the receiver operating curve (AUROC) with a 90% confidence interval obtained by 1000x bootstrapping. Pink = best, yellow = second-best method. For RCC subtyping, AUROCs from top to bottom refer to clear cell, chromophobe and papillary RCC. Statistical significance is reported in **Suppl. Tables 3-7**.

|  | Renal Cell Ca.<br>subtype<br>AACHEN<br>N=248 | Colorectal<br>MSI<br>TCGA<br>N=426 | Colorectal<br>BRAF<br>TCGA<br>N=500 | Gastric<br>MSI<br>TCGA<br>N=327 | Gastric<br>EBV<br>TCGA<br>N=327 | Bladder<br>FGFR3<br>TCGA<br>N=241 |
| --- | --- | --- | --- | --- | --- | --- |
|  | 0.955<br>(0.929 - 0.978) |  |  |  |  |  |
| ResNet | 0.964<br>(0.935 - 0.988) | 0.867<br>(0.821 - 0.908) | **0.795**<br>**(0.739 - 0.848)** | 0.68<br>(0.611 - 0.746) | 0.764<br>(0.674 - 0.852) | 0.782<br>(0.704 - 0.849) |
|  | 0.952<br>(0.928 - 0.982) |  |  |  |  |  |
|  | 0.969<br>(0.952 - 0.982) |  |  |  |  |  |
| EfficientNet | 0.908<br>(0.819 - 0.972) | 0.764<br>(0.702 - 0.818) | 0.744<br>(0.677 - 0.802) | **0.742**<br>**(0.685 - 0.801)** | 0.629<br>(0.554 - 0.703) | 0.746<br>(0.683 - 0.809) |
|  | 0.945<br>(0.915 - 0.969) |  |  |  |  |  |
|  | 0.973<br>(0.958 - 0.985) |  |  |  |  |  |
| ViT | 0.971<br>(0.929 - 0.998) | **0.907**<br>**(0.873 - 0.941)** | 0.781<br>(0.719 - 0.834) | 0.73<br>(0.671 - 0.789) | 0.721<br>(0.619 - 0.820) | **0.785**<br>**(0.719 - 0.84)** |
|  | 0.97<br>(0.952 - 0.984) |  |  |  |  |  |
|  | 0.961<br>(0.942 - 0.977) |  |  |  |  |  |
| MIL | 0.969<br>(0.951 - 0.984) | 0.586<br>(0.523 - 0.647) | 0.559<br>(0.492 - 0.624) | 0.498<br>(0.433 - 0.565) | **0.789**<br>**(0.715 - 0.857)** | 0.584<br>(0.501 - 0.662) |
|  | 0.912<br>(0.875 - 0.948) |  |  |  |  |  |
|  | 0.964<br>(0.945 - 0.980) |  |  |  |  |  |
| CLAM | 0.957<br>(0.905 - 0.994) | 0.614<br>(0.545 - 0.682) | 0.622<br>(0.553 - 0.688) | 0.712<br>(0.650 - 0.776) | 0.727<br>(0.637 - 0.808) | 0.644<br>(0.542 - 0.74) |
|  | 0.96<br>(0.936 - 0.981) |  |  |  |  |  |

**Figure 2.**
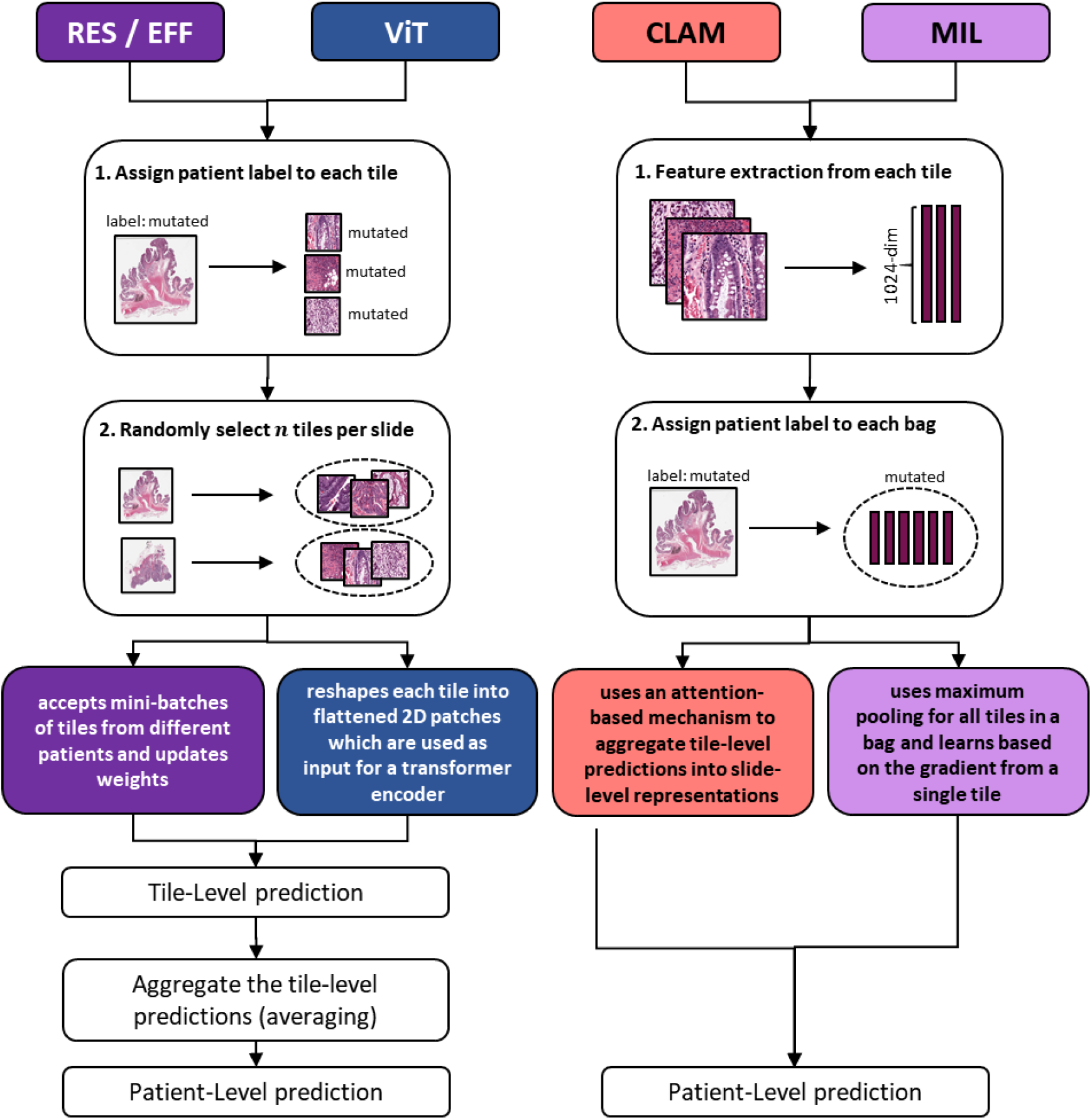
Schematic workflow of the methods. ResNet (RES) and EfficientNet (EFF) as well as Vision Transformers (ViT) were used for weakly supervised end-to-end prediction benchmark tasks. In addition, clustering-constrained attention multiple-instance learning (CLAM) and classical multiple instance learning (MIL) were used for the same tasks. While classical workflows use different models (RES, EFF, ViT), they all cast slide labels to image tiles. In contrast, CLAM and classical MIL cast slide labels to bags of image tiles without assuming that every single tile reflects the target of interest.

**Figure 3.**
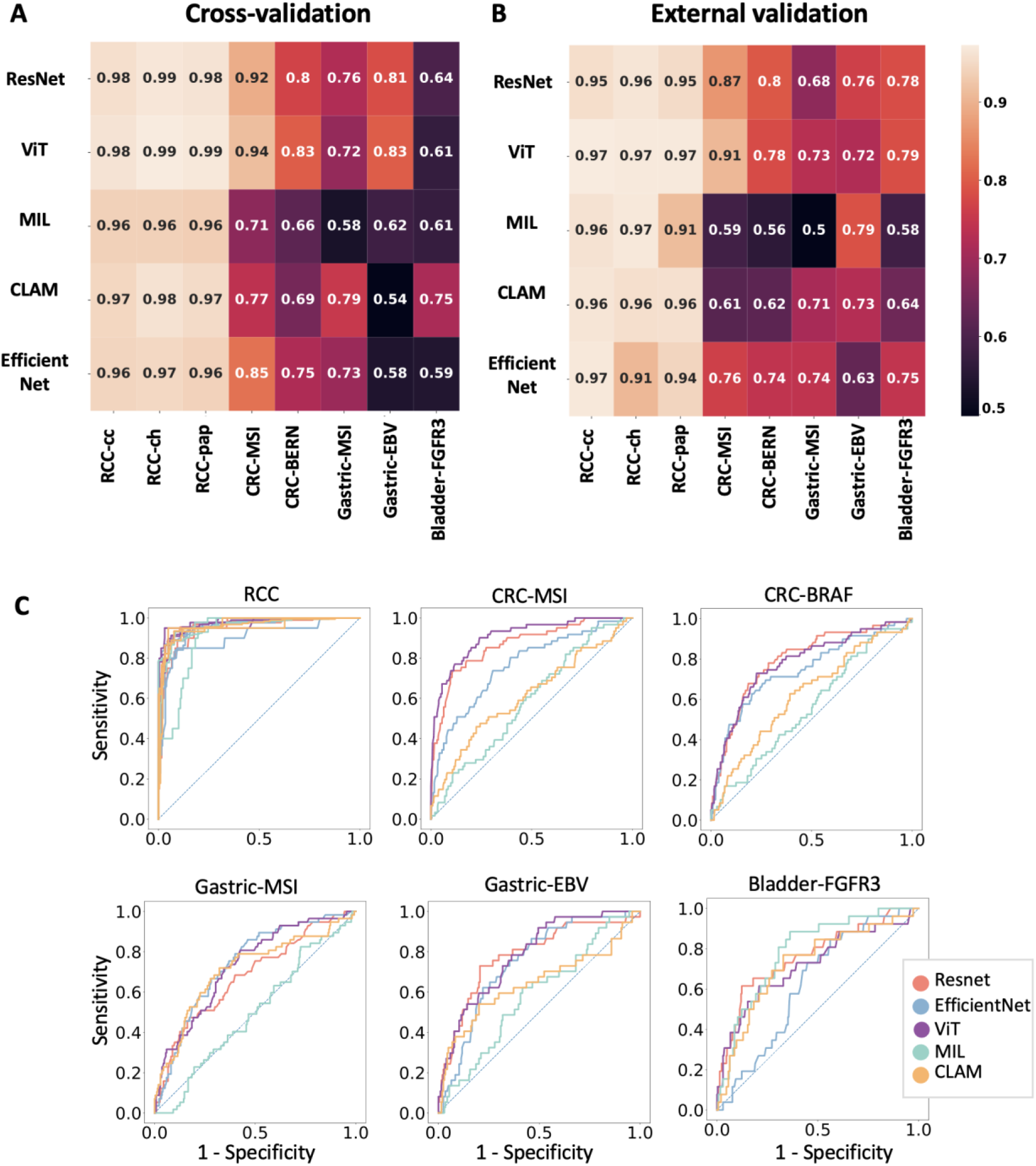
Benchmarking results. A) Cross-validation area under the receiver operating curve (AUROC) for each method. B) External validation AUROC, C) Receiver operating curves show that RCC subtyping is almost perfectly solved by all methods while molecular subtyping tasks are solved best by ViT and ResNet.

### ViT and ResNet excel in mutation prediction in colorectal cancer

Next, we focused on prediction of clinically actionable genetic alterations directly from H&E histology WSI: MSI and BRAF in colorectal cancer, MSI and EBV in gastric cancer and FGFR3 mutations in bladder cancer. In a cross-validated experiment in the large DACHS cohort of colorectal cancer, ViT achieved a state-of-the art AUROC of 0.937 (0.919 -0.953; N=2039 patients). The classical ResNet-based approach achieved the second-highest performance with an AUROC of 0.919 (0.899 -0.937). Both classifiers generalized well to the external validation cohort (TCGA-CRC, N=426 patients) with ViT and ResNet again yielding the highest and second-highest performance for MSI prediction with AUROCs of 0.907 (0.873 -0.941) and 0.867 (0.821 -0.908), respectively. Compared to the other approaches EfficientNet, MIL and CLAM, the performance was significantly higher (z>4 and p<0.0001 for all, **Suppl. Table 5**). Although ViT slightly outperformed ResNet (z=1.17), the direct comparison ViT and ResNet did not reach statistical significance (p=0.24, **Suppl. Table 3**). All other methods, in particular MIL-based methods reached much lower performances in within-cohort experiments, with classical MIL and CLAM yielding AUROC of 0.715 (0.657 -0.77) and 0.771 (0.716 -0.825), respectively (**Table 1**). Likewise, in external validation experiments, MIL and CLAM yielded the lowest performance (**Table 2**) which was statistically significantly inferior to all other approaches (p<=0.01, **Suppl. Table 6 and 7**). Prediction of BRAF mutational status (N=2075 patients in cross-validation) resulted in the same ranking of algorithms with ViT achieving the highest (AUROC 0.83 [0.799 -0.858]), ResNet the second-highest (AUROC 0.801 [0.772 -0.831]) and classical MIL achieving the lowest performance (AUROC 0.661 [0.592 -0.727]). Also in external validation (**Table 2**), ResNet (AUROC 0.795 [0.739 -0.848]) and ViT (0.781 [0.719 -0.834]) significantly (p<=0.02) outperformed all other approaches (**Suppl. Table 3** and **Suppl. Table 5**).

### Prediction of molecular alterations in gastric and bladder cancer

While colorectal cancer is among the most widely studied tumor types in computational pathology, it is important to validate computational methods also in rarer tumor types. [7] Therefore, we tested all five algorithms on prediction of the clinically relevant alterations MSI and Epstein-Barr Virus (EBV) in gastric cancer and FGFR3 mutations in bladder cancer. We found that the overall performance in our proprietary datasets (BERN for gastric, AACHEN for bladder cancer) was lower than for colorectal cancer, which is in line with previous studies. [17,18] The highest AUROCs were 0.785 (0.715 -0.852) for MSI in gastric cancer (N=302 patients), 0.831 (0.68 -0.957) for EBV in gastric cancer (N=304 patients) and 0.748 (0.636 -0.85) for FGFR3 in bladder cancer (N=87 patients, **Table 1**). The highest performance was achieved by CLAM in gastric MSI and bladder FGFR3 and by ViT in gastric EBV, while the second-highest performance was always achieved by the classical ResNet-based workflow. In the external validation experiment for gastric cancer (TCGA-STAD with N=327 patients for MSI, N=327 patients for EBV), the resulting performance differences were much less clear-cut (**Table 2**), with no consistently best-performing method. However, for external validation of FGFR3 analysis in bladder cancer (TCGA-BLCA, N=241 patients), ViT and ResNet again outperformed all other approaches, reaching AUROCs of 0.785 (0.719 -0.84) and 0.782 (0.704 -0.849), respectively. The difference between ResNet and MIL and CLAM was statistically significant (p<=0.03, Suppl. Table 3).

### Overall assessment of classifier performance for mutation prediction

Finally, we systematically analyzed performance differences between the five classifiers in all five mutation prediction tasks. Each method was compared to the four other methods in five tasks, yielding 20 comparisons per method. ResNet significantly (p<0.03, z>2) outperformed other methods in 8/20 tasks and was never significantly outperformed by another method (**Suppl. Table 3**). ViT significantly (p<=0.002, z>3) outperformed other methods in 7/20 tasks and was never significantly outperformed (**Suppl. Table 5**). EfficientNet outperformed other methods in 6/20 tasks, but was outperformed in 3/20 tasks (**Suppl. Table 4**). MIL outperformed other methods in 1/20 tasks but was outperformed in 13/20 tasks (**Suppl. Table 6**). Similarly, CLAM outperformed other methods in 1/20 mutation prediction tasks but was outperformed in 7/20 tasks (**Suppl. Table 7**). Overall, we conclude that ViT and ResNet-based approaches are reasonable algorithm choices for prediction of molecular alterations from routine histology in solid tumors.

### Explainability of the performance differences

To understand the reason for the observed performance differences of the methods, we systematically compared which image tiles were assigned the highest scores by each method, in all classification tasks. We found that for renal cell carcinoma subtyping -a task in which all methods performed almost equally well -highest scoring tiles showed plausible histopathological patterns for all classes for all methods. Consistently, tiles with high prediction scores for clear cell RCC showed carcinoma cells with clear cytoplasm; tiles predictive of chromophobe RCC showed a perinuclear halo characteristic of this subtype and tiles with high scores for papillary RCC showed a papillary tissue architecture (**Figure 4**). In contrast, for MSI prediction in colorectal cancer -a task in which classical end-to-end methods outperformed MIL-based methods -the typical MSI-like morphology [19] includes poor differentiation, mucinous differentiation and tumor-infiltrating lymphocytes. These patterns were prominently visible in highly scoring tiles selected by high-performing methods ResNet, EfficientNet and ViT. In contrast, MIL-based methods assigned the highest prediction scores to image tiles at the tissue boundary, less than half of which clearly showed MSI-like morphology (**Figure 5**). We conclude that the performance of end-to-end AI methods is directly related to the ability to assign high prediction scores to image tiles with informative histopathological patterns and thus, performance is directly linked to histopathological phenotype.

**Figure 4.**
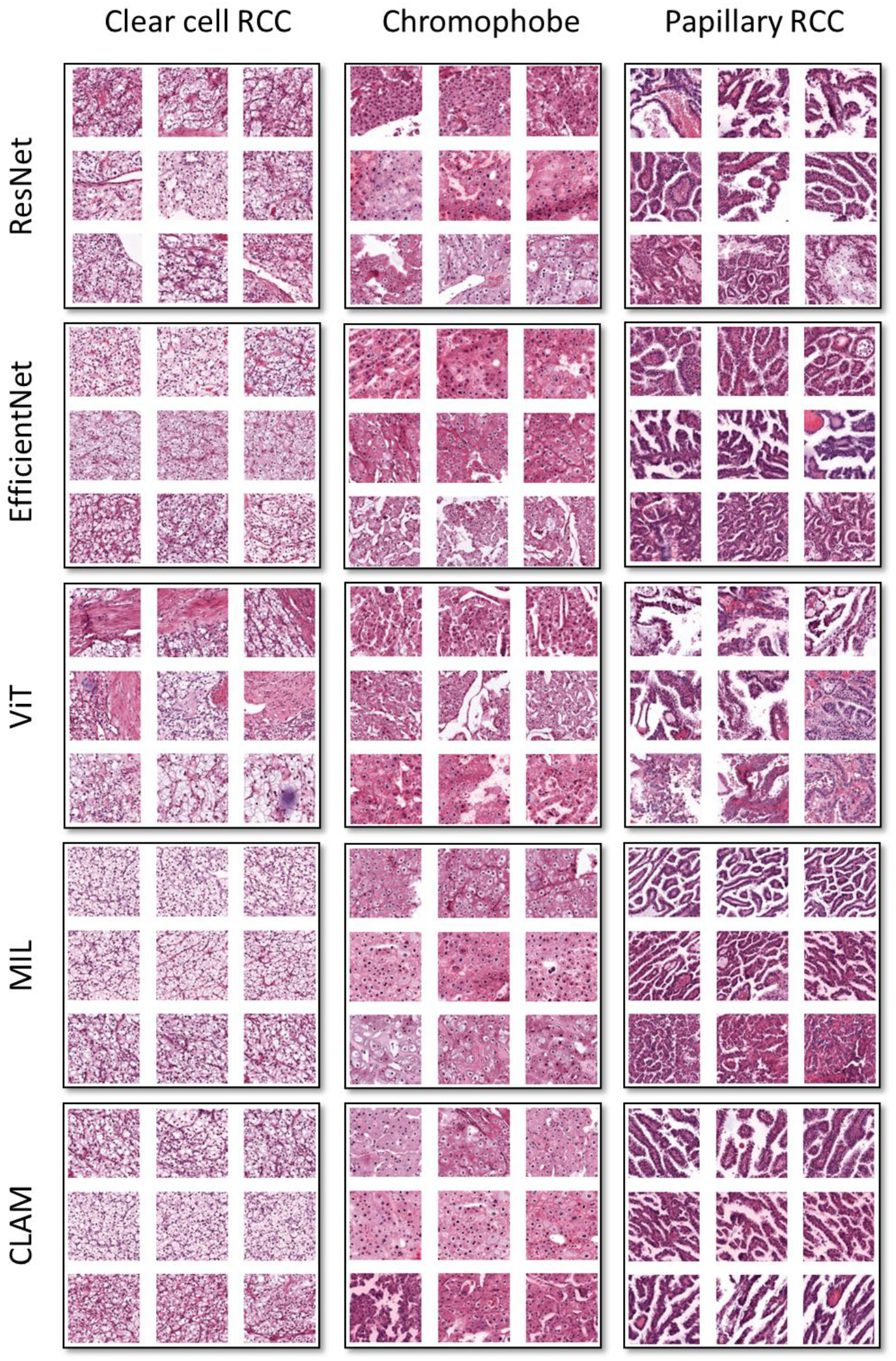
Explainability of subtyping of renal cell carcinoma (RCC). The three highest scoring tiles for the three highest scoring patients in the external validation experiment as selected by each method are displayed. For this benchmark task, all six methods achieved a high performance. Correspondingly, all methods succeeded in selecting image tiles with patterns representative of known features of RCC subtypes.

**Figure 5.**
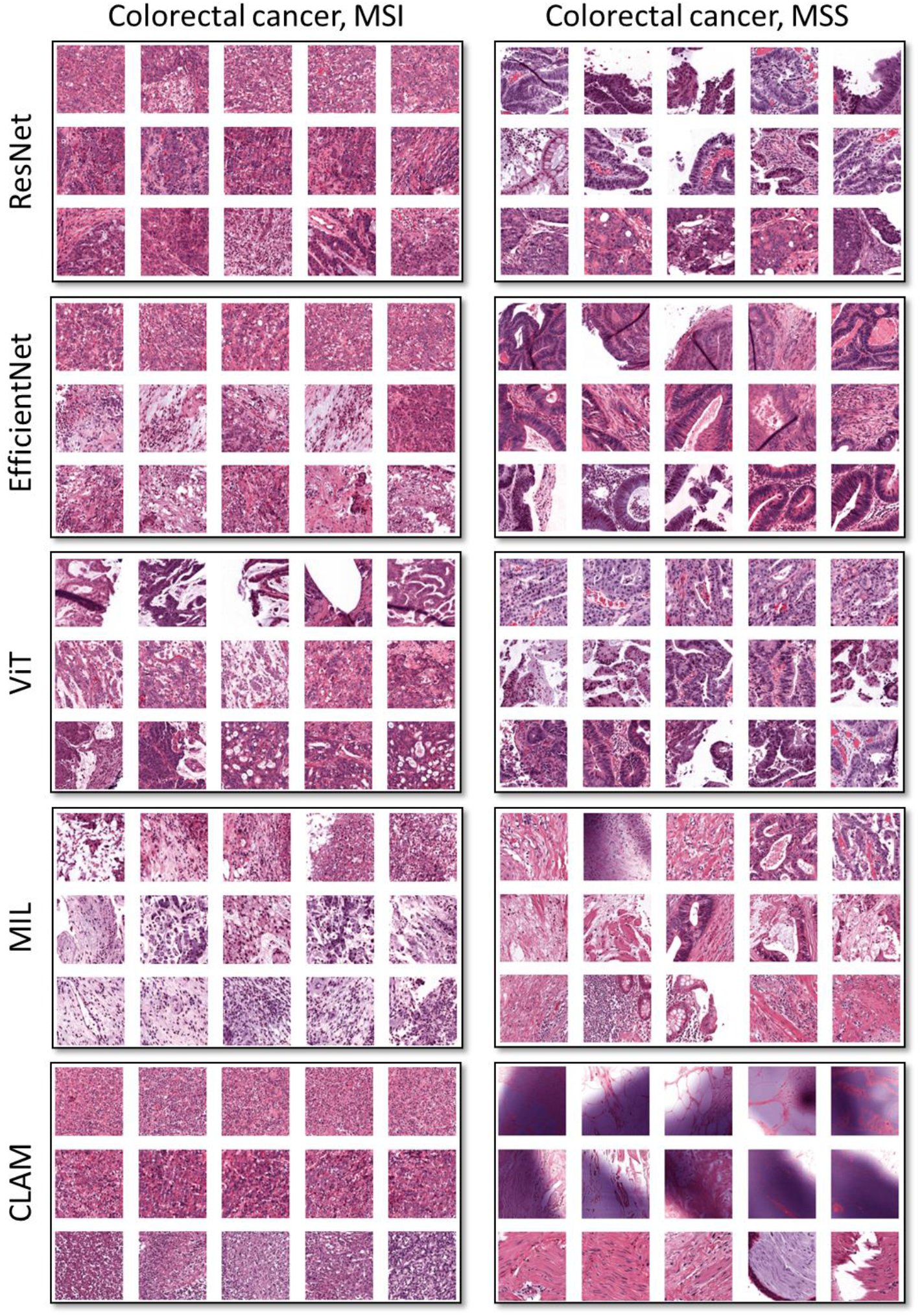
Explainability of microsatellite instability (MSI) prediction in colorectal cancer (CRC). The five highest scoring tiles for the three highest scoring patients in the external validation experiment are displayed. Resnet, EfficientNet and ViT achieved the highest performance. This corresponds to a selection of biologically plausible tiles, showing poorly differentiated, mucinous tumors for MSI. Conversely, MIL and CLAM selected tiles with tissue edges and other artifacts, corresponding to their poor performance.

## Discussion

In this study, we provide a systematic benchmark for five AI algorithms applied to six clinical problems in computational pathology. We chose these particular six problems because they were previously addressed in one or several publications and are of direct diagnostic or therapeutic relevance. [2,5,17,18,20] We demonstrate that morphological subtyping of renal cell carcinoma (RCC) is an easy task which can be solved by any common computational pathology method with high performance (**Table 1** and **Table 2**), without significant differences between methods (**Suppl. Tables 3, 4, 5, 6 and 7**).

However, prediction of clinically targetable molecular alterations directly from histology uncovered pronounced differences between different approaches: Overall, classical weakly supervised workflows (in which all tiles inherit the slide label) markedly outperformed MIL-based workflows (in which labels are only defined for bags of tiles). While classical approaches are being used since 2018 [10,18], MIL has been first used in a large-scale computational pathology study in 2019 [1] Because the classical MIL is highly susceptible to artifacts and classifier instability, the newer MIL-based variant CLAM has been shown to be more robust and powerful than classical MIL. [2] CLAM performs well for morphological subtyping of lung cancer and renal cell carcinoma [2] as well as for prediction of primary tumor type from metastatic tissue [2,12] However, our data demonstrate that researchers should choose classical weakly supervised workflows rather than MIL-based workflows for mutation prediction tasks. Although the classical end-to-end approaches investigated in this study suffer from label noise (they assign the label to all tiles generated from a slide, not just the tumor tissue), this does not seem to impair performance when a large portion of the slide is tumor, as in the surgical resection specimen in this study. It is possible that such label noise will lead to a lower performance in needle-in-a-haystack problems such as detection of small nests of tumor cells in biopsy tissue [1] or in lymph nodes. [21] Another possibility for lower performance of MIL/CLAM than the classical weakly supervised approach is that a pre-trained network was used for feature extraction in MIL/CLAM whereas networks were trained directly on images in the classical weakly supervised approach. However, during training of the classical weakly supervised approach, we only trained the deepest 50% of all layers, essentially using shallow network layers as a pre-trained feature extractor. Also, previous studies have demonstrated that pre-trained features can in principle be used for high-performing mutation prediction in cancer.[22] In summary, our benchmark study provides important actionable advice for future studies and real-world applications of computational pathology, in particular by showing that label noise does not impair performance for mutation prediction tasks on surgical resection tissue.

Within weakly supervised workflows, ResNet or ViT are the model architectures of choice. ResNet is the de facto standard in computational pathology because of its high efficiency with comparatively few parameters. ViT performed on par with but never significantly outperformed ResNet (**Suppl. Table 5**). This finding is of high practical relevance for academic and commercial actors in computational pathology, as ViTs represent a relatively novel technology, which has been broadly applied outside of medicine but is still new to computational pathology. More generally, these data show that new AI approaches which were established in non-medical fields of research can be applied relatively easily to the domain of pathology, provided that clinically relevant benchmark tasks are analyzed.

There are multiple limitations of our study: it is in the nature of technical benchmarks that neither all possible technical approaches nor all possible applications can be evaluated. In this study, we selected five technical approaches and six previously studied applications of clinical relevance. In a recent systematic analysis [7], we found that classical weakly-supervised workflows and MIL-based approaches account for almost all deep learning studies in tumor subtyping and prediction of molecular alterations. Regarding the clinical applications, we investigate molecular subtyping in colorectal, gastric and bladder cancer, i.e. a very common and two less common tumor types. In addition, our colorectal cancer cohorts comprised more patients than the gastric and bladder cancer cohort (**Suppl. Table 1**). Importantly, as part of our study we release an open-source workflow that includes all five approaches: the histology image analysis package (HIA). HIA is a comprehensive PyTorch-based library which enables academic and commercial researchers to easily benchmark all tested methods on their own datasets, using just a single implementation. HIA allows to apply all methods to other clinical tasks and also extend the toolkit by plugging in new classifiers, but using the existing pre/post-processing pipeline. We expect that our findings and this tool can help computational pathology to reach clinical-grade performance and ultimately have a positive impact on treatment selection and resource saving in the healthcare system.

## Methods

### Ethics statement and patient cohorts

All experiments were conducted in accordance with the Declaration of Helsinki. For this study we used anonymized H&E stained slides obtained from formalin-fixed paraffin-embedded (FFPE) material from the “The Cancer Genome Atlas” (TCGA) archive (available at https://portal.gdc.cancer.gov), a large, multi-centric collection of tissue specimen obtained from multiple hospitals across different countries. From this cohort, we only used “diagnostic slides”, i.e. digitized images of glass slides which were used by the respective medical center to make the diagnosis of cancer. In addition, we used four proprietary datasets: the DACHS study (“Darmkrebs: Chancen der Verhütung durch Screening”), a large population-based case-control and patient cohort study on CRC, including samples of patients with stages I-IV from different laboratories in southwestern Germany coordinated by the German Cancer Research Center (Heidelberg, Germany) [23,24]. The DACHS study was approved by the ethics committees of the University of Heidelberg and of the Medical Chambers of Baden-Württemberg and Rhineland-Palatinate, and all participants signed an informed consent. The BERN dataset is a single-center dataset collected from clinical routine samples at the pathology archive at Inselspital, University of Bern (Bern, Switzerland) [25]. Use of this data set was approved by the local ethics commission, specifically granting the use of archival tissue for molecular and immunohistochemical analysis as well as tissue microarray construction (University of Bern, Switzerland, no. 200/14). The use of archival tissue from this cohort for molecular analysis was approved by the local ethical commission (Technical University of Munich, No. 2136/08). Similarly, the AACHEN-RCC dataset and the AACHEN-BLADDER datasets originated from a single high-volume medical center, the pathology archive at RWTH Aachen University Hospital (Aachen, Germany). The collection of patient samples from Aachen was approved by the local Ethics board (AACHEN-RCC: EK315/19, AACHEN-BLADDER: EK455/20). All cohorts were anonymized at the time of analysis. **Suppl. Table 1** shows patient numbers and a clinico-pathological description of all cohorts.

### Prediction tasks and experimental design

In this study, we benchmarked technical approaches in six end-to-end prediction tasks, i.e. we trained AI algorithms to predict each of these targets from raw histological whole slide images: (1) Diagnosis of renal cell carcinoma subtype (clear cell RCC, chromophobe RCC and Papillary RCC); (2) prediction of microsatellite instability (MSI) or mismatch repair deficiency (dMMR) in colorectal cancer; (3) prediction of BRAF mutation in colorectal cancer; (4) prediction of microsatellite instability (MSI) or mismatch repair deficiency (dMMR) in gastric cancer; (5) detection of Epstein-Barr Virus (EBV) presence in gastric cancer and (6) prediction of FGFR3 point mutations in bladder cancer. MSI and dMMR have a very high degree of overlap and are interchangeably used in clinical routine. [26] Here, we use the term “MSI” throughout the study.

We pre-defined the following experimental design: First, for each prediction task, we used one cohort for within-cohort experiments by patient-level three-fold cross-validation. For this experiment, we used the following cohorts: DACHS-CRC, BERN-Gastric, Aachen-Bladder, TCGA-RCC. Subsequently, we re-trained a classifier for each prediction task on the training cohorts and externally validated it in a separate patient cohort. For external validation, we used the following cohorts: TCGA-CRC, TCGA-Gastric, TCGA-Bladder, Aachen-RCC. The validation cohorts were not used for any other purpose except for validation of the final model. We did not perform any hyperparameter tuning but used a pre-defined set of hyperparameters for each method (**Table 3**).

**Table 3:** Hyperparameters and technical details for all approaches.

|  | Hyperparameters and architecture | Reference, technical | Reference, medical |
| --- | --- | --- | --- |
| ResNet | ● Resnet-18 pre-trained on ImageNet (18 layers, the last layer was changed from 1000 output neurons to N output neurons for N classes)<br>● Epochs = 8 (batch size = 128)<br>● Maximum number of tiles = 512<br>● Optimizer = Adam<br>● learning rate = 1e-4 (weight decay = 1e-5)<br>● Freeze ratio of layers =0.5 (weights and biases in the first 9 layers were not trainable, weights and biases in the last 9 layers were trainable) | [34] | [5,22] |
| EfficientNet | ● Pre-trained on ImageNet (efficientnet-b7)<br>● Epochs = 8 (batch size = 128)<br>● Maximum number of tiles = 512<br>● Optimizer = Adam<br>● learning rate = 1e-4 (weight decay = 1e-5)<br>● Freeze ratio of layers =0.5 | [42] | [43] |
| ViT | ● Pre-trained on ImageNet (B_32_imagenet1k, 24 layers, the last layer was changed from 1000 output neurons to N output neurons for N classes)<br>● Epochs = 8 (batch size = 128)<br>● Maximum number of tiles = 512<br>● Optimizer = Adam<br>● learning rate = 1e-4 (weight decay = 1e-5)<br>● Freeze ratio of layers =0.5 | [14,44] | N/A |
| MIL | ● Extract features using a Resnet-50 (which is pre-trained on ImageNet) for all the tiles of a slide<br>● Epochs = 50 (batch size = 1)<br>● Optimizer = Adam<br>● learning rate = 1e-4 (weight decay = 1e-5)<br>● dropout = True | [45] | [1] |
| CLAM | ● Extract features using a Resnet-50 (which is pre-trained on ImageNet) for all the tiles of a slide<br>● Epochs = 50 (batch size = 1)<br>● Optimizer = Adam<br>● learning rate = 1e-4 (weight decay = 1e-5)<br>● Model size = small<br>● Bag loss = Cross Entropy<br>● Instance Loss = SmoothTop1SVM[46]<br>● dropout = True | [2] | [2] |

### Ground truth for prediction tasks

The ground truth for the prediction targets were obtained as follows: For TCGA-CRC and TCGA-STAD, MSI and EBV status were obtained from a public source [27] as described before [5]. For TCGA-RCC, images from the three morphological subtypes were obtained separately from the GDC data portal (TCGA-KIRP for papillary, KIRC for clear cell and KICH for chromophobe tumors). In DACHS, MSI status was obtained by 3-plex PCR and BRAF V600E mutational status was obtained by immunohistochemistry on tissue microarrays and by Sanger sequencing as described before [28,29] For the BERN gastric cancer cohort, MSI/dMMR status was obtained with immunohistochemistry for DNA repair enzymes and EBV status was obtained by Epstein-Barr virus (EBV)-encoded RNA (EBER) in-situ hybridization. AACHEN-BLADDER comprised bladder carcinomas from a real-world cohort [17] and FGFR3 mutational status was obtained by whole exome sequencing or identified using the SNaPshot method. [30] In the AACHEN-RCC cohort, morphological subtype was retrieved from the routine pathology report.

### Image preprocessing

The input images for all the methods were preprocessed based on the “Aachen protocol for Deep Learning histopathology” [31]. Based on this protocol, the digitized whole slide images were tessellated into smaller image tiles of (512 ×512) pixels at a resolution of 0.5 micrometers per pixel (MPP). During this process, tiles containing background and artifacts were removed from the data set (using canny edge detection in Python’s OpenCV package). Extracted tiles were color normalized using the Macenko method to reduce the inter-cohort color bias [32]. No manual annotations were applied to the whole slide images and all subsequent AI methods were trained exclusively with slide-level labels.

### Artificial intelligence methods

For our benchmarking task, we implemented and systematically compared five different methods for end-to-end artificial intelligence on WSI (**Table 3**).

For “classical” methods, we followed a workflow established previously for lung cancer [10] and colorectal cancer [5,18]. Briefly, this algorithm is based on the assumption that all tiles from a given slide inherit the slide label for classification. AI algorithms are trained on N randomly selected tiles per WSI and tile-level predictions are averaged to obtain patient-level predictions. WSIs contain tumor and non-tumor tissue and only tumor tissue and tumor-adjacent normal tissue is expected to reflect the target label. However, empirically, such weakly supervised methods can yield clinical-grade performance despite weak labels. [15,33] Three different AI models were used within this classical approach: ResNet, EfficientNet and Vision Transformers (ViT).

1. ResNets are currently the de-facto standard for supervised transfer learning due to their higher performance and efficiency when compared to other CNN models [34]. In this study, we used a ResNet18 model with only the 50% deepest layers as trainable layers. The model was pre-trained on ImageNet and fine-tuned by transfer learning on each benchmark task separately.

2. EfficientNet aims to scale up the baseline convolutional network which has been referred to as EfficientNet-B0 [35]. The common approach in designing any ConvNet is to develop a smaller version of the network and then scale it up to reach the higher performance. EfficientNet scales the width, depth and resolution of the network using the compound scaling method, which achieves state-of-the-art accuracies on smaller and therefore faster networks.

3. ViT is the most modern AI architecture analyzed within the classical workflows. Since 2017, attention-based models have become the dominant selection in natural language processing (NLP) [36]. In 2020, a high performance of transformers in visual tasks has been demonstrated [14]. The input to the vision transformer are flattened 2D patches extracted from the original image. All the layers of the transformer use a constant latent vector size (D). Through a patch embedding block, the flattened patches get mapped to D dimensions using a trainable linear projection. This step is followed by a position embedding block which adds positional information to each patch. The encoder of the transformer consists of alternating layers of self-attention, multilayer perceptron (MLP), layer norm (LN) before each block and residual connections after each block. Although ViTs showed very good performance on ImageNet data set, its performance on histopathological images with smaller size has not been systematically investigated before this study.

The conceptual limitations of the classical weakly-supervised computational pathology workflow are addressed by multiple instance learning (MIL). MIL does not simply cast the slide label on each tile, but rather groups tiles in “bags”. While the label of instances is not clear for the model, the label of the bag is positive, if there is at least one positive instance within that bag. Otherwise, the label of the bag would be negative, if all the instances are negative. Thus, MIL is in theory well suited to handle a heterogenous set of tiles obtained from different regions in a WSI. In this study, we tested two established MIL methods.

4. Classical MIL has been used diversely in processing of histopathological images due to the lack of annotation for the tiles extracted from whole slide images [37–40] The basic framework of MIL is creating bags containing different number of instances and was in the past successfully applied to large-scale image classification tasks in histopathology [1]

5. Clustering constrained Attention Multiple instance learning (CLAM) has been designed initially to overcome the challenges in the standard MIL approaches [2]. By using attention-based deep learning methods, it is able to detect the most informative regions on a WSI which was empirically shown to outperform classical MIL in some classification tasks [2]. Compared to standard MIL methods, which use the gradient signal only from one single instance from each bag to update the learning parameters, CLAM aggregates patch-level features into slide-level information required for classification, thus achieving higher robustness. CLAM uses low-dimensional features extracted from the input tiles (which is computationally expensive), but the actual training only uses feature vectors and the required computational power and time for training of this model is very low. The source code for CLAM and MIL methods are taken from https://github.com/mahmoodlab/CLAM and were modified based on our workflow. Figure 2 shows the workflow for each model.

### Statistics

The primary statistical endpoint was the area under the receiver operating curve (AUROC) calculated on the level of patients. Confidence intervals were obtained by 1000 bootstrapping the AUC computation. For this purpose, we sampled with replacement from the original ground truth labels and the predictions and recomputed the new AUC value. 90% confidence interval is selected from the sorted AUC values. For binary classification tasks, AUROCs were identical for both groups and therefore, only the AUROC for the positive group (mutated, MSI/dMMR) is reported. For multiclass classification tasks, we binarized the ground truth labels (for each class) and calculated the AUROC for the prediction scores of the same class (Macro-averaging). To quantify whether performance differences between models were statistically significant, we used DeLong’s method. This method tests whether two models have a significant difference in their performance and accounts for the role of randomness in the finite datasets [41]. The output of this method is the z score (difference of AUROC of the output performance of two models divided by its standard error) and the p value.

### Code availability

All methods are implemented using Python 3.8 with PyTorch and all source codes for preprocessing are available at https://github.com/KatherLab/preProcessing and all codes for training and evaluating the models with the Histology Image Analysis package are available at https://github.com/KatherLab/HIA under an open source license.

## Funding

JNK is supported by the German Federal Ministry of Health (DEEP LIVER, ZMVI1-2520DAT111) and the Max-Eder-Programme of the German Cancer Aid (grant #70113864). CT is supported by the German Research Foundation (DFG) (SFB CRC1382, SFB-TRR57). The DACHS study was supported by the German Research Council (BR 1704/6-1, BR 1704/6-3, BR 1704/6-4, CH 117/1-1, HO 5117/2-1, HO 5117/2-2, HE 5998/2-1, KL 2354/3-1, RO 2270/8-1 and BR 1704/17-1); the Interdisciplinary Research Program of the National Center for Tumor Diseases (NCT), Germany; and the German Federal Ministry of Education and Research (01KH0404, 01ER0814, 01ER0815, 01ER1505A, and 01ER1505B). PB is supported by the DFG, German Research Foundation (Project-IDs 322900939, 454024652, 432698239, 445703531, 445703531), European Research Council (ERC; Consolidator Grant AIM.imaging.CKD, No 101001791), Federal Ministry of Education and Research(STOP-FSGS-01GM1901A), and Federal Ministry of Economic Affairs and Energy (EMPAIA, No. 01MK2002A).NTG is funded by the DFG (GA 1384/3-1, GA 1384/5-1).

## Disclosures

JNK declares consulting services for Owkin, France and Panakeia, UK. TJB reports owning a company that develops mobile apps, outside the scope of the submitted work (Smart Health Heidelberg GmbH, Handschuhsheimer Landstr. 9/1, 69120 Heidelberg). No other potential conflicts of interest are reported by any of the authors.

---

**Suppl. Table 1:** Clinico-pathological features of all cohorts. N/A not applicable.

|  | **TCGA**<br>**CRC** | **DACHS**<br>**CRC** | **TCGA**<br>**STAD** | **BERN**<br>**STAD** | **TCGA**<br>**BLCA** | **AC**<br>**BLCA** | **TCGA**<br>**RCC** | **AC**<br>**RCC** |
| --- | --- | --- | --- | --- | --- | --- | --- | --- |
| **N total** | 500 | 2244 | 327 | 304 | 241 | 87 | 897 | 248 |
| **female** | 245<br>(49%) | 931<br>(42%) | 107<br>(33%) | 110<br>(36%) | 57<br>(24%) | 42<br>(48%) | N/A | N/A |
| **male** | 253<br>(51%) | 1313<br>(59%) | 220<br>(67%) | 194<br>(64%) | 184<br>(76%) | 45<br>(52%) | N/A | N/A |
| **Stage I** | 71<br>(14%) | 406<br>(18%) | 42<br>(13%) | 35<br>(12%) | 0<br>(0%) | N/A | N/A | N/A |
| **Stage II** | 152<br>(30%) | 765<br>(34%) | 101<br>(31%) | 41<br>(14%) | 80<br>(33%) | N/A | N/A | N/A |
| **Stage III** | 134<br>(27%) | 757<br>(34%) | 150<br>(46%) | 116<br>(38%) | 82<br>(34%) | N/A | N/A | N/A |
| **Stage IV** | 58<br>(12%) | 315<br>(14%) | 32<br>(10%) | 1 (0.33%) | 77<br>(32%) | N/A | N/A | N/A |
| **MSI** | 61<br>(12%) | 210<br>(9%) | 57<br>(17%) | 42<br>(14%) | N/A | N/A | N/A | N/A |
| **MSS** | 365<br>(73%) | 1829<br>(82%) | 270<br>(83%) | 260<br>(86%) | N/A | N/A | N/A | N/A |
| **BRAF m** | 59<br>(12%) | 151<br>(7%) | N/A | N/A | N/A | N/A | N/A | N/A |
| **BRAF**<br>**wt** | 441<br>(88%) | 1924<br>(86%) | N/A | N/A | N/A | N/A | N/A | N/A |
| **EBV+** | N/A | N/A | 26<br>(8%) | 8<br>(3%) | N/A | N/A | N/A | N/A |
| **EBV-** | N/A | N/A | 301<br>(92%) | 296<br>(97%) | N/A | N/A | N/A | N/A |
| **FGFR3**<br>**m** | N/A | N/A | N/A | N/A | 3(15%) | 13<br>(15%) | N/A | N/A |
| **FGFR3**<br>**wt** | N/A | N/A | N/A | N/A | 204<br>(85%) | 74<br>(85%) | N/A | N/A |
| **subtype**<br>**KIRP** | N/A | N/A | N/A | N/A | N/A | N/A | 275<br>(31%) | 44<br>(18%) |
| **subtype**<br>**KIRC** | N/A | N/A | N/A | N/A | N/A | N/A | 512<br>(57%) | 184<br>(74%) |
| **subtype**<br>**KICH** | N/A | N/A | N/A | N/A | N/A | N/A | 109<br>(12.15%) | 20<br>(8.06%) |
| **Ref.** | [47] | [48] | [49] | [25] | [50] | [17] | [51] | N/A |

**Suppl. Table 2:** Area under the precision recall curve (AUPRC) for all external validation experiments.

|  |  | **Renal Cell Ca.**<br>**subtype**<br>**AACHEN**<br>**N=248** | **Colorectal**<br>**MSI**<br>**TCGA**<br>**N=426** | **Colorectal**<br>**BRAF**<br>**TCGA**<br>**N=500** | **Gastric**<br>**MSI**<br>**TCGA**<br>**N=327** | **Gastric**<br>**EBV**<br>**TCGA**<br>**N=327** | **Bladder**<br>**FGFR3**<br>**TCGA**<br>**N=241** |
| --- | --- | --- | --- | --- | --- | --- | --- |
|  |  | 0.982<br>(0.969 - 0.993) | 0.615<br>(0.502 - 0.718) | 0.362<br>(0.268 - 0.474) | 0.310<br>(0.229 - 0.409) | 0.294<br>(0.164 - 0.451) | 0.436<br>(0.303 - 0.564) |
|  | **ResNet** | 0.727<br>(0.548 - 0.874) |  |  |  |  |  |
|  |  | 0.886<br>(0.807 - 0.957) |  |  |  |  |  |
|  |  | 0.989<br>(0.984 - 0.994) | 0.397<br>(0.292 - 0.520) | 0.311<br>(0.229 - 0.409) | 0.342<br>(0.260 - 0.452) | 0.101<br>(0.07 - 0.148) | 0.280<br>(0.198 - 0.385) |
|  | **EfficientNet** | 0.713<br>(0.525 - 0.844) |  |  |  |  |  |
|  |  | 0.792<br>(0.682 - 0.871) |  |  |  |  |  |
|  |  | 0.991<br>(0.986 - 0.995) | 0.712<br>(0.622 - 0.795) | 0.377<br>80.28 - 0.476) | 0.362<br>(0.266 - 0.477) | 0.293<br>(0.159 - 0.446) | 0.433<br>(0.289 - 0.563) |
|  | **ViT** | 0.885<br>(0.753 - 0.983) |  |  |  |  |  |
|  |  | 0.879<br>(0.804 - 0.938) |  |  |  |  |  |
|  |  | 0.987<br>(0.980 - 0.993) | 0.193<br>(0.14 - 0.257) | 0.174<br>(0.115 - 0.245) | 0.164<br>(0.128 - 0.207) | 0.219<br>(0.139 - 0.380) | 0.185<br>(0.134 - 0.266) |
|  | **MIL** | 0.575<br>(0.397 - 0.735) |  |  |  |  |  |
|  |  | 0.877<br>(0.801 - 0.94) |  |  |  |  |  |
|  |  | 0.988<br>(0.981 - 0.994) | 0.282<br>(0.199 - 0.382) | 0.171<br>(0.126 - 0.225) | 0.403<br>(0.287 - 0.502) | 0.171<br>(0.107 - 0.274) | 0.306<br>(0.202 - 0.442) |
|  | **CLAM** | 0.840<br>(0.699 - 0.943) |  |  |  |  |  |
|  |  | 0.835<br>(0.741 - 0.919) |  |  |  |  |  |

**Suppl. Table 3:** Pairwise comparison of classifier performance, relative to ResNet. Z scores and p values were obtained with DeLong’s test. p<0.05 was considered statistically significant and all respective cells are highlighted (yellow: ResNet is significantly better, blue: ResNet is significantly worse compared to the reference method).

|  |  | **EfficientNet** | **ViT** | **MIL** | **CLAM** |
| --- | --- | --- | --- | --- | --- |
| morpho-<br>logical<br>subtyping | **ResNet**<br>**RCC-cc** | z = -0.82<br>p = 0.41 | z = -1.02<br>p = 0.31 | z = -0.34<br>p = 0.73 | z = -0.49<br>p = 0.62 |
|  | **ResNet**<br>**RCC-ch** | z = 0.90<br>p = 0.37 | z = -0.59<br>p = 0.56 | z = 1.13<br>p = 0.26 | z = -0.13<br>p = 0.89 |
|  | **ResNet**<br>**RCC-pap** | z = 0.85<br>p = 0.39 | z = -0.32<br>p = 0.75 | z = -0.28<br>p = 0.78 | z = 0.19<br>p = 0.85 |
| predicting<br>molecular<br>alterations | **ResNet**<br>**CRC-MSI** | z = 2.27<br>p = 0.02 | z = -1.17<br>p = 0.24 | z = 5.93<br>p < 0.0001 | z = 5.24<br>p < 0.0001 |
|  | **ResNet**<br>**CRC-BRAF** | z = 0.97<br>p = 0.33 | z = 0.29<br>p = 0.77 | z = 4.24<br>p < 0.0001 | z = 3.24<br>p = 0.001 |
|  | **ResNet**<br>**Gastric-MSI** | z = -1.16<br>p = 0.24 | z = -0.99<br>p = 0.32 | z = 3.36<br>p = 0.0007 | z = -0.54<br>p = 0.59 |
|  | **ResNet**<br>**Gastric-EBV** | z = 1.86<br>p = 0.06 | z = 0.58<br>p = 0.56 | z = 0.340<br>p = 0.69 | z = 0.51<br>p = 0.61 |
|  | **ResNet**<br>**Bladder-**<br>**FGFR3** | z = 0.56<br>p = 0.58 | z = -0.05<br>p = 0.96 | z = 3.20<br>p = 0.001 | z = 2.11<br>p = 0.03 |

**Suppl. Table 4:** Pairwise comparison of classifier performance, relative to EfficientNet. Z scores and p values were obtained with DeLong’s test. p<0.05 was considered statistically significant and all respective cells are highlighted (yellow: EfficientNet is significantly better, blue: EfficientNet is significantly worse compared to the reference method).

|  |  | **ResNet** | **ViT** | **MIL** | **CLAM** |
| --- | --- | --- | --- | --- | --- |
| morpho-<br>logical<br>subtyping | **EfficientNet**<br>**RCC-cc** | z = 0.82<br>p = 0.41 | z = -0.32<br>p = 0.74 | z = 0.64<br>p = 0.52 | z = 0.40<br>p = 0.69 |
|  | **EfficientNet**<br>**RCC-ch** | z = -0.90<br>p = 0.37 | z = -1.20<br>p = 0.23 | z = -0.08<br>p = 0.93 | z = -0.85<br>p = 0.39 |
|  | **EfficientNet**<br>**RCC-pap** | z = -0.85<br>p = 0.39 | z = -1.27<br>p = 0.20 | z = -1.19<br>p = 0.23 | z = -0.67<br>p = 0.50 |
| predicting<br>molecular<br>alterations | **EfficientNet**<br>**CRC-MSI** | z = -2.27<br>p = 0.02 | z = -4.20<br>p < 0.0001 | z = 3.54<br>p = 0.0004 | z = 2.66<br>p = 0.007 |
|  | **EfficientNet**<br>**CRC-BRAF** | z = -0.97<br>p = 0.33 | z = -0.76<br>p = 0.45 | z = 3.41<br>p = 0.0006 | z = 2.56<br>p = 0.01 |
|  | **EfficientNet**<br>**Gastric-MSI** | z = 1.16<br>p = 0.24 | z = 0.22<br>p = 0.82 | z = 4.98<br>p < 0.0001 | z = 0.51<br>p = 0.61 |
|  | **EfficientNet**<br>**Gastric-EBV** | z = -1.86<br>p = 0.06 | z = -1.32<br>p = 0.19 | z = -2.38<br>p = 0.02 | z = -1.31<br>p = 0.19 |
|  | **EfficientNet**<br>**Bladder-**<br>**FGFR3** | z = -0.56<br>p = 0.58 | z = -0.80<br>p = 0.42 | z = 2.27<br>p = 0.02 | z = 1.47<br>p = 0.14 |

**Suppl. Table 5:**
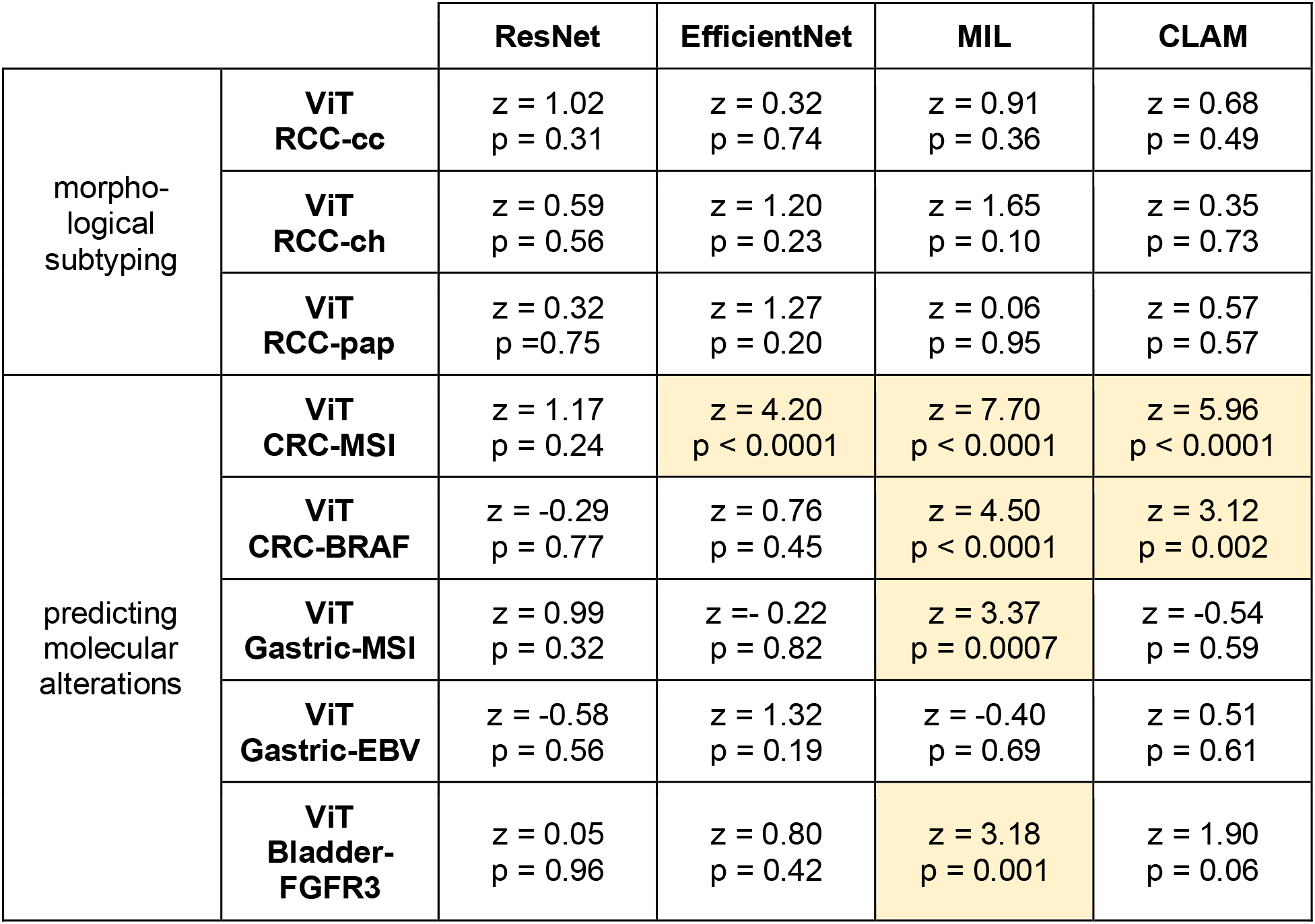
Pairwise comparison of classifier performance, relative to ViT. Z scores and p values were obtained with DeLong’s test. p<0.05 was considered statistically significant and all respective cells are highlighted (yellow: ViT is significantly better, blue: ViT is significantly worse compared to the reference method).

**Suppl. Table 6:** Pairwise comparison of classifier performance, relative to MIL. Z scores and p values were obtained with DeLong’s test. p<0.05 was considered statistically significant and all respective cells are highlighted (yellow: MIL is significantly better, blue: MIL is significantly worse compared to the reference method).

|  |  | **ResNet** | **EfficientNet** | **ViT** | **CLAM** |
| --- | --- | --- | --- | --- | --- |
| morpho-<br>logical<br>subtyping | **MIL**<br>**RCC-cc** | z = 0.34<br>p = 0.73 | z = -0.64<br>p = 0.52 | z = -0.91<br>p = 0.36 | z = -0.18<br>p = 0.85 |
|  | **MIL**<br>**RCC-ch** | z = -1.13<br>p = 0.26 | z = 0.08<br>p = 0.93 | z = -1.65<br>p = 0.10 | z = -1.27<br>p = 0.20 |
|  | **MIL**<br>**RCC-pap** | z = 0.28<br>p = 0.78 | z = 1.19<br>p = 0.23 | z = -0.06<br>p = 0.95 | z = 0.55<br>p = 0.58 |
| predicting<br>molecular<br>alterations | **MIL**<br>**CRC-MSI** | z = -5.93<br>p < 0.0001 | z = -3.54<br>p = 0.0004 | z = -7.70<br>p < 0.0001 | z = -0.49<br>p = 0.62 |
|  | **MIL**<br>**CRC-BRAF** | z = -4.24<br>p < 0.0001 | z = -3.41<br>p = 0.0006 | z = -4.50<br>p < 0.0001 | z = -1.17<br>p = 0.24 |
|  | **MIL**<br>**Gastric-MSI** | z = -3.36<br>p = 0.0007 | z = -4.98<br>p < 0.0001 | z = -3.37<br>p = 0.0007 | z = -3.43<br>p < 0.0001 |
|  | **MIL**<br>**Gastric-EBV** | z = -0.340<br>p = 0.69 | z = 2.38<br>p = 0.02 | z = 0.40<br>p = 0.69 | z = 0.96<br>p = 0.34 |
|  | **MIL**<br>**Bladder-**<br>**FGFR3** | z = -3.20<br>p = 0.001 | z = -2.27<br>p = 0.02 | z = -3.18<br>p = 0.001 | z = -0.79<br>p = 0.43 |

**Suppl. Table 7:** Pairwise comparison of classifier performance, relative to CLAM. Z scores and p values were obtained with DeLong’s test. p<0.05 was considered statistically significant and all respective cells are highlighted (yellow: CLAM is significantly better, blue: CLAM is significantly worse compared to the reference method).

|  |  | ResNet | EfficientNet | ViT | MIL |
| --- | --- | --- | --- | --- | --- |
| morpho-<br>logical<br>subtyping | **CLAM**<br>**RCC-cc** | z = 0.49<br>p = 0.62 | z = -0.40<br>p = 0.69 | z = -0.68<br>p = 0.49 | z = 0.18<br>p = 0.85 |
|  | **CLAM**<br>**RCC-ch** | z = 0.13<br>p = 0.89 | z = 0.85<br>p = 0.39 | z = -0.35<br>p = 0.73 | z = 1.27<br>p = 0.20 |
|  | **CLAM**<br>**RCC-pap** | z = -0.19<br>p = 0.85 | z = 0.67<br>p = 0.50 | z = -0.57<br>p = 0.57 | z = -0.55<br>p = 0.58 |
| predicting<br>molecular<br>alterations | **CLAM**<br>**CRC-MSI** | z = -5.24<br>p < 0.0001 | z = -2.66<br>p = 0.007 | z = -5.96<br>p < 0.0001 | z = 0.49<br>p = 0.62 |
|  | **CLAM**<br>**CRC-BRAF** | z = -3.24<br>p = 0.001 | z = -2.56<br>p = 0.01 | z = -3.12<br>p = 0.002 | z = 1.17<br>p = 0.24 |
|  | **CLAM**<br>**Gastric-MSI** | z = 0.54<br>p = 0.59 | z = -0.51<br>p = 0.61 | z = 0.54<br>p = 0.59 | z = 3.43<br>p < 0.0001 |
|  | **CLAM**<br>**Gastric-EBV** | z = -0.51<br>p = 0.61 | z = 1.31<br>p = 0.19 | z = -0.51<br>p = 0.61 | z = -0.96<br>p = 0.34 |
|  | **CLAM**<br>**Bladder-**<br>**FGFR3** | z = -2.11<br>p = 0.03 | z = -1.47<br>p = 0.14 | z = -1.90<br>p = 0.06 | z = 0.79<br>p = 0.43 |

